# Cysteine String Protein alpha in Extracellular Vesicle Subtypes: a Proteomic Analysis

**DOI:** 10.1101/2023.12.13.571333

**Authors:** Luiz Gustavo Nogueira de Almeida, Victoria Armstrong, Antoine Dufour, Janice E.A. Braun

## Abstract

Cysteine string protein (CSPα /DnaJC5) is a presynaptic J-domain protein (JDP) that prevents neurodegeneration. CSPα/DnaJC5 is reported to facilitate export of distinct, highly oligomeric, disease-causing proteins in addition to wild-type TDP-43, tau and α-synuclein. Yet, detailed mechanistic knowledge of the full CSPα/DnaJC5 secreted proteome is lacking. Understanding the CSPα/DnaJC5 export pathway has implications for a growing number of neurodegenerative diseases. In humans, Leu115Arg or Leu116deletion mutations cause adult-onset neuronal ceroid lipofusinosis (ANCL), a rare neurodegenerative disorder. In the present study, we examined extracelular vesicles (EVs) released from CSPα/DnaJC5 expressing cells. Cells are known to secrete many types of EVs of different sizes and origins into the extracellular space. EV subpopulations were separated by their sedimentation speed and subjected to proteomic analysis. We find that CSPα/DnaJC5 and the CSPα/DnaJC5 mutants, Leu115Arg or Leu116del are enriched in multiple EV subpopulations. The exported protein profile is determined by proteomics. We report that several other J-domain proteins (JDPs), such as DnaJC7, DnaJA1 and DnaJA2 are exported and speculate that export of JDPs may facilitate the secretion of diverse client proteins. Our work provides a platform for further inquiry into the role of secreted CSPα/DnaJC5 and other JDPs in proteostasis.

## Introduction

An extensive network of cellular machinery tightly regulates the synthesis, function and elimination of proteins. In addition to the autophagy/lysosomal and ubiquitin/proteasomal pathways of protein degradation, cells can rid themselves of toxic and misfolded proteins by secreting them. Cysteine string protein (CSPα/DnaJC5), a protein encoded by the DnaJC5 gene, is an evolutionary conserved neuroprotective J-domain containing protein (JDP) that ejects proteins (Braun and Scheller, 1995, Deng et al., 2017). Removal of proteins via CSPα/DnaJC5 is distinct from the import of proteins into lysosomes by chaperone-mediated autophagy (Lee et al., 2018). In the absence of CSPα/DnaJC5, mice, *drosophila* and *C. elegans* undergo rapid neurodegeneration, underscoring the essential nature of CSPα/DnaJC5 export in the maintenance of functional neurons (Zinsmaier et al., 1994, Fernandez-Chacon et al., 2004, Kashyap et al., 2014). The evidence supporting CSPα/DnaJC5-mediated export of proteins is substantial (Wu et al., 2023, Deng et al., 2017, Fontaine et al., 2016, Lee et al., 2018, Pink et al., 2021). CSPα/DnaJC5 is found in cell-derived lipid bilayer enclosed particles, called extracellular vesicles (EVs) (Deng et al., 2017, Pink et al., 2021). While some EVs are directly formed and released from the plasma membrane, other vesicles are first generated through endosomal, multivesicular bodies that subsequently fuse with the plasma membrane and release their internal vesicles into the extracellular milieu (Pegtel and Gould, 2019, Cheng and Hill, 2022).

JDPs are a molecular chaperone family with 50 members in humans that carry out diverse functions (Kampinga and Craig, 2010, Kampinga et al., 2009, Zhang et al., 2022, Cyr and Ramos, 2023). CSPα/DnaJC5 protein export is one of many JDP-regulated pathways. The JDPs, DnaJC23/Sec63 and DnaJC19/Tim14 are essential in protein translocation into the endoplasmic reticulum and mitochondria, respectively (Itskanov et al., 2021, Mokranjac et al., 2006). DnaJC6/auxillin facilitates the uncoating of clathrin from clathrin coated vesicles (Eisenberg and Greene, 2007), DnaJC13/RME8 is involved in endosomal trafficking, DnaJB6 ensures quality control of nuclear pore complexes (Kuiper et al., 2022) and DnaJC29/sacsin is implicated in cytoskeleton dynamics, organelle positioning and synaptic adhesion (Francis et al., 2022, Girard and McPherson, 2008, Romano et al., 2022). DnaJB11/ERdj3 is an endoplasmic reticulum JDP that transits the classical secretory pathway to enter the extracellular space and bind extracellular proteins (Genereux et al., 2015). In contrast, DnaJB1 is exported in EVs, enhancing the proteostasis capacity of cells that receive DnaJB1-containing EVs (Takeuchi et al., 2015). As well, some JDPs are implicated in viral life cycles (Taguwa et al., 2015, Chand et al., 2023). Each JDP contains a J domain with a conserved histidine, proline, aspartate (HPD) motif that intracellularly accelerates ATPase of Hsp70s (HSPA) through a transient JDP-Hsp70 interaction (Kampinga and Craig, 2010). Several JDPs are involved in managing neurodegenerative-disease-related proteins and mutations in JDPs are linked to multiple heritable diseases (Ayala Mariscal and Kirstein, 2021, Rosenzweig et al., 2019, Zhang et al., 2022, Zarouchlioti et al., 2018).

CSPα/DnaJC5 contains an N-terminal J domain and a middle region displaying a string of 13–15 cysteine residues that are extensively palmitoylated (Braun and Scheller, 1995). There is a small N-terminal segment proceeding the J domain and the cysteine residues are followed by a presumably unstructured C-terminal (Figure 1A). The structure of the J domain has been solved by nuclear magnetic resonance (Patel et al., 2016). The role of post-translational modifications in regulating CSPα/DnaJC5 is well-established. CSPα/DnaJC5 is phosphorylated at serine 10 and 34 (Evans et al., 2001, Evans and Morgan, 2005, Patel et al., 2016, Shirafuji et al., 2018) and extensively palmitoylated within the cysteine string region (Gundersen et al., 1994, Braun and Scheller, 1995). Phosphorylation alters CSPα/DnaJC5’s association with client proteins (Evans and Morgan, 2002) while palmitoylation influences self-association and dissociation as well as cellular localization (Swayne et al., 2003, van de Goor and Kelly, 1996, Chamberlain and Burgoyne, 1998, Arnold et al., 2004, Greaves and Chamberlain, 2006, Greaves et al., 2008, Stowers and Isacoff, 2007, Ohyama et al., 2007, Braun and Scheller, 1995, Xu et al., 2010, Greaves et al., 2012, Zhang and Chandra, 2014, Diez-Ardanuy et al., 2017). The major palmitoylating and depalmitoylating enzymes for CSPα/DnaJC5 are HIP14/DHHC17 and PPT1, respectively (Stowers and Isacoff, 2007, Ohyama et al., 2007).

**Figure 1.**
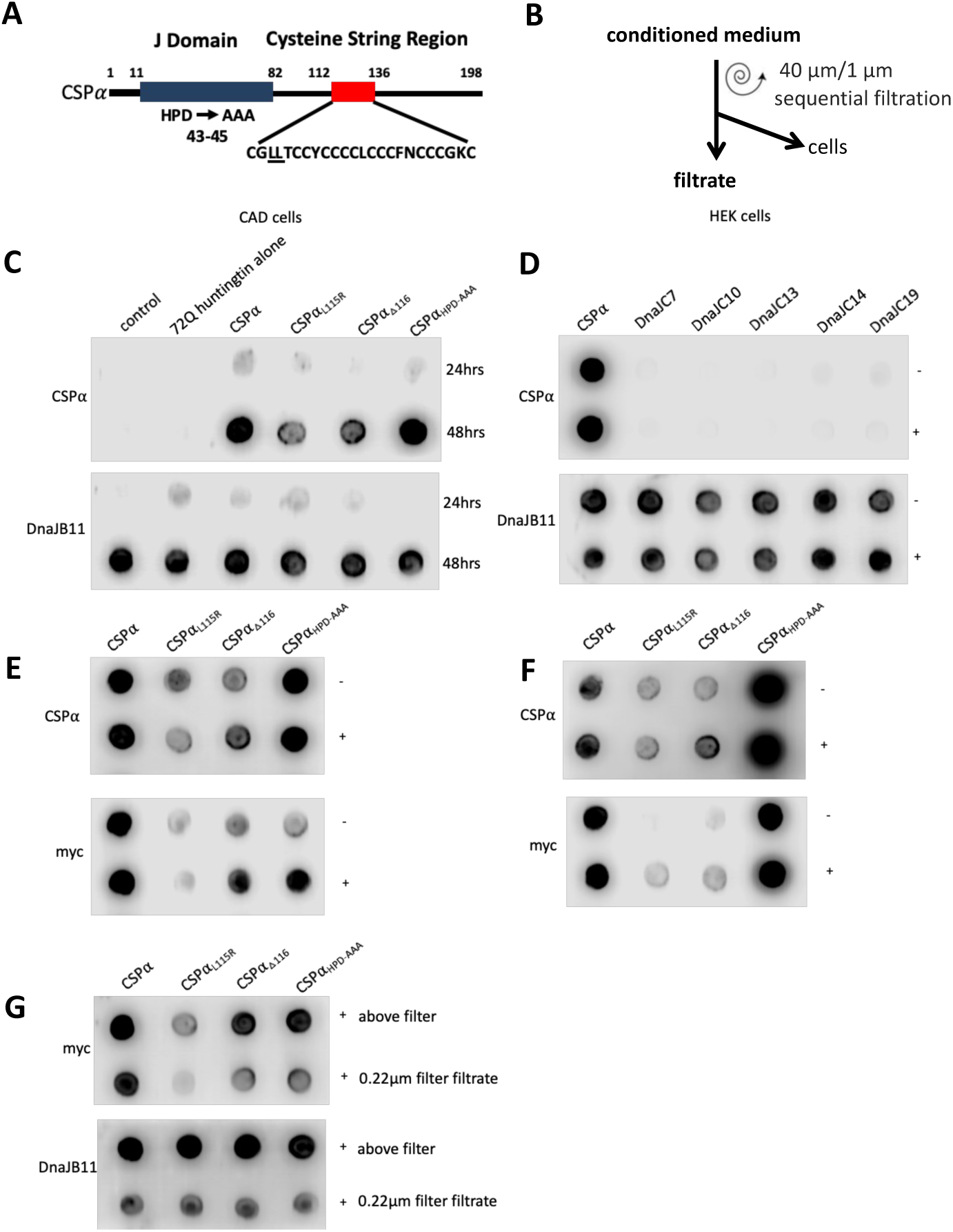
CSPα secretion from CAD and HEK293 cells. (A) Schematic diagram of CSPα. Domains are highlighted and ANCL-causing-mutations are indicated. (B) Scheme of unfractionated conditioned medium collection. (C-G) Dot Blots are probed for CSPα, DnaJB11 and myc as indicated. (C) Media from control CAD cells, and cells expressing GFP-72Q huntingtin^exon1^ or CSPα, CSPα_L115R_, CSPα_L116del_, and CSPα_HPD-AAA_ in the presence of GFP-72Q huntingtin^exon1^ 24 h and 48 h after transfection. (D) Media from HEK cells expressing the indicated JDPs in the absence (-) or presence (+) of GFP-72Q huntingtin_exon1_ 48 h after transfection. (E) Media from CAD cells expressing myc-tagged CSPα, CSPα_L115R_, CSPα_L116del_, and CSPα_HPD-AAA_ co-transfected in the absence (-) or presence (+) of GFP-72Q huntingtin_exon1_ 48 h after transfection. (F) Media from HEK cells co-transfected with CSPα, CSPα_L115R_, CSPα_L116del_, and CSPα_HPD-AAA_ in the absence (-) or presence (+) of GFP-72Q huntingtin^exon1^ 48 h after transfection. (G) Media was further filtered through millex-GV 0.22 µm PVDF filters.

CSPα/DnaJC5 exports wild-type and mutant α-synuclein, TDP-43 and tau from HEK293 cells as well as mutant SOD-1 and misfolded huntingtin from CAD cells (Fontaine et al., 2016, Deng et al., 2017). Soluble proteins as well as proteins enclosed within EVs are exported. In the presence of CSPα/DnaJC5, GFP-72Q huntingtin^exon1^ is exported in two EV subtypes as determined by microflow cytometry (180-240nm) and widefield microscopy (10-30µm) (Pink et al., 2021). In contrast, CSPα/DnaJC5 facilitates secretion of wild-type and A30P, E46K and A53T α-synuclein mutants as soluble species not enclosed within EVs from HEK293 and COS7 cells (Wu et al., 2023, Lee et al., 2018). Remarkably, ExoU, a potent toxin injected into cells by *Pseudomonas aeruginosa* is delivered to the plasma membrane in host cells by CSPα/DnaJC5 (Deruelle et al., 2021). ExoU is a phospholipase A2 that causes cell lysis, highlighting the range of proteins transported by the CSPα/DnaJC5 pathway. In addition, to the removal of cellular proteins CSPα/DnaJC5 is reported to modulate ciliogenesis (Piette et al., 2021), but how this relates mechanistically to the the export of α-synuclein, TDP-43, tau, SOD-1 and huntingtin requires further study.

Together, these studies show that CSPα/DnaJC5 is a JDP that facilitates synapse maintenance and that interfering with CSPα/DnaJC5 function causes synapse instability and leads to neurodegeneration. In addition, CSPα/DnaJC5 plays a prominent role in the cellular export of heterogeneous protein cargo without common structural motifs via EVs and/or as non-vesicular protein aggregates.

Despite these substantial advances many questions remain. Mechanistic details of how CSPα/DnaJC5 mutations lead to neurotoxicity are not yet established. Detailed molecular knowledge of EVs that contain CSPα/DnaJC5 is lacking. This question is important because we lack the understanding of whether the CSPα/DnaJC5 pathway is an endogenous protein export pathway that is hijacked by toxic disease-causing-proteins proteins or if CSPα/DnaJC5 is solely responsible for balancing proteostasis when misfolded protein production exceeds autophagy/lysosomal and ubiquitin/proteasomal elimination capacity. Here we undertook a proteomic approach to identify the full CSPα/DnaJC5 exported proteome utilizing a cell-based secretion assay. The protein composition of EVs pelleted at high (100K), medium (10K) and low (2K) speeds were compared and the validated exported proteins determined. Our proteomics screen expands the repertoire of known CSPα/DnaJC5 exported proteins as well as the repertoire of exported JDPs in EVs.

## Results and Discussion

### CSPα/DnaJC5 secretion from CAD and HEK293 cells

We first evaluated the classical and unconventional export of JDPs from CAD and HEK293 cell lines transiently transfected with CSPα/DnaJC5 by monitoring CSPα/DnaJC5 and DnaJB11/ERdj3 in unfractionated media (Figure 1). While CSPα/DnaJC5 undergoes unconventional secretion, DnaJB11/ERdj3 has an N-terminus signal peptide and travels through the classical secretory pathway (Genereux et al., 2015). Conditioned media was collected from transfected cells grown without serum for 24 h and 48 h, filtered and 10 µL spotted on nitrocellulose and evaluated. Expression of CSPα/DnaJC5 in CAD or HEK293 cells increases extracellular CSPα/DnaJC5 protein levels in conditioned media. The ANCL mutants, CSPα_L115R_, CSPα_L116del_ are secreted as is CSPα_HPD-AAA_ (Figure 1C). Endogenous DnaJB11/ERdj3 secretion is observed at 48 h in the absence and presence of CSPα, CSPα mutants, and several other JDPs (Figure 1C, D). Since CSPα/DnaJC5 exports different types of insoluble proteins, we next co-expressed CSPα/DnaJC5 together with GFP-72Q huntingtin_exon1_ and examined export. CSPα and CSPα mutants were secreted in the presence of misfolded GFP-72Q huntingtin_exon1_ in both CAD and HEK293 cells (Figure 1D, E, F). It is noteworthy that less CSPα_L115R_ and CSPα_L116del_ export was observed in comparison to CSPα/DnaJC5 and CSPα_HPD-AAA._ When the media was filtered through 0.22 µm poly-vinylidene fluoride (PVDF) hydrophillic filters, CSPα/DnaJC5 and DnaJB11 were detected in the filtrate as well as above the filter indicating these JDPs were associated with extracellular entities of different sizes (Figure 1G). Thus, we confirm that CSPα/DnaJC5, CSPα_L115R_, CSPα_L116del_ and CSPα_HPD-AAA_ are exported in both the presence and absence of aggregation-prone proteins.

### Proteomic profiling of extracellular subtypes secreted from CSPα/DnaJC5 expressing cells

Using our established export assay, we sought to determine the CSPα/DnaJC5 secreted proteome in the absence of expressed misfolded proteins. To do so, we further separated conditioned media from control and CSPα/DnaJC5 expressing CAD cells by differential ultracentrifugation and performed extensive proteomic analysis by in-solution liquid chromatography with tandem mass spectrometry (LC-MS/MS). Pelleted materials were recovered by low (2,000 x *g* = 2K) or medium (10,000 x *g* = 10K) or high (100,000 x *g* = 100K) centrifugation of media collected from cells. The 10K and 100K pellets were then washed, recentrifuged and resuspended (Figure 2A). Proteome analysis was conducted on conditioned media collected from cells grown without serum addition (n = 3) and in medium supplemented with 7.5% exosome-depleted serum (n = 3). This workflow allowed for comparisons of exported proteins from control and CSPα/DnaJC5 expressing cells without introducing the EVs routinely present in serum. CSPα/DnaJC5 expression reduced cell viability (Figure 2B). The total amount of protein pelleted at 2K was higher than the 10K and 100K pellets, as expected, and the total protein was highest in the 2K pellet from CSPα/DnaJC5 expressing cells (Figure 2C). Greater autofluorescence of the 2K pellet from CSPα/DnaJC5-expressing cells was observed (Figure 2D & Supplementary Figure 1). In order to evaluate exported debris the 2K pellet was not washed. The 2K pelleted fractions include exported debris and does not differentiate between adhering proteins, large secreted membrane vesicles and secreted protein aggregates. We compared the distribution of CSPα/DnaJC5 and DnaJB11/ERdj3 in the 2K, 10K and 100K pellets by dot blot (Figure 2E). CSPα/DnaJC5 was found in all three pelleted fractions. As anticipated, EVs isolated by centrifugation are heterogeneous with the majority of smaller EVs in the 100K pellet, classically considered exosomes (Figure 2F). 100 µg of each fraction was analyzed by mass spectrometry. Because exported proteins were identified from 100 µg of the corresponding pelleted fractions the actual higher protein content of the 2K pellet obtained from CSPα/DnaJC5 expressing cells is not reflected in the proteomics results. Standard EV markers were detectable in EVs pelleted at high speed as well as in the larger/heavier EVs pelleted at medium or low speed. Release of EVs is dynamic and serum starvation as well as CSPα/DnaJC5 expression led to changes in EV markers. Yet, despite these changes, CD81 and programmed cell death 6-interacting protein (PDCD6IP or ALIX) were among the most abundant EV markers identified in EVs pelleted at high speed (Figure 2G).

**Figure 2.**
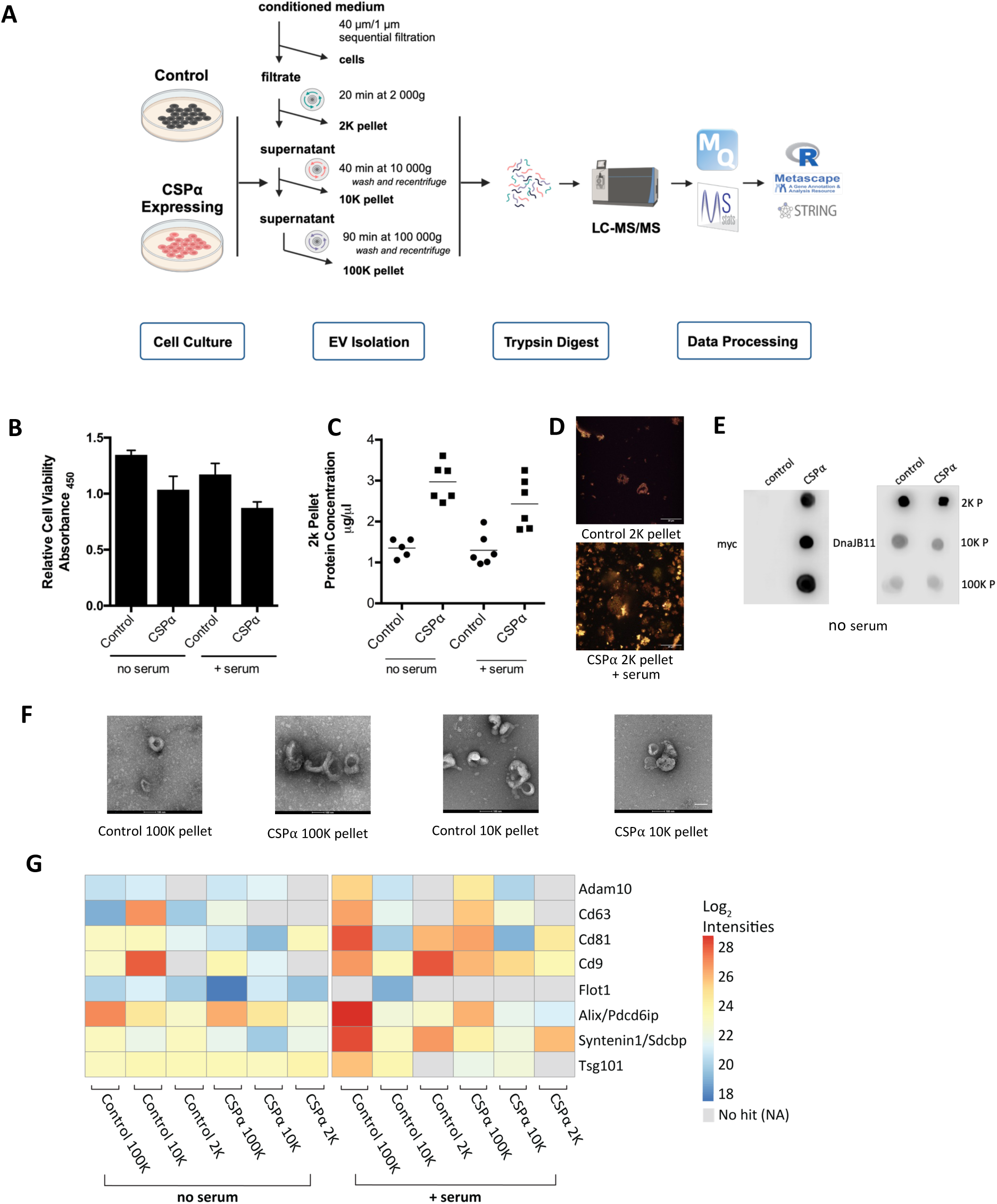
Heterogeneous EVs released from control and CSPα-expressing cells. (A) Work flow of 2K, 10K and 100K pellets from conditioned media generated from CAD cells expressing CSPα/DnaJC5. (B) Cell viability in media with no serum and media with 7.5% exosome-free FBS. (C) Protein content of the 2K pellet. (D) Representative image of 2K pellet autofluorescence. (E) Dot blot validation of the indicated fractions probed for CSPα/DnaJC5 and DnaJB11/ERdj3 (F) Representative images of 10K and 100K pellet fractions. (G) Heat map of classical EV markers. The heat map shows proteins that were independently identified in 2 (out of 3) biological samples.

A continuum of pelleted fractions containing CSPα/DnaJC5 was observed indicating CSPα/DnaJC5 is present in a range of EVs that are separated by their pelleting properties (Figure 3A). Low levels of CSPα/DnaJC5 export were detected in control cells by LC-MS/MS. Given that JDPs have an indespensible role in proteostasis, we scrutinized the possible export of other JDPs. 22 distinct JDPs, in addition to CSPα/DnaJC5, were found to be exported from CAD cells. Importantly, this estimate is conservative in that only JDPs that were independently identified in 2 out of 3 biological samples are shown and were considered significant. The 2K pellets from cells stressed by serum starvation contained the largest number of JDPs (Figure 3). Some JDPs were exported at a higher level, for example DnaJC7, DnaJA1 and DnaJA2 were found in all three pelleted fractions from both serum fed and serum starved cells (Figure 3). Their export occurred in the presence and absence of CSPα/DnaJC5 expression. DnaJC7, DnaJA1 and DnaJA2 do not show homology with CSPα/DnaJC5 outside of the J domain but they have been linked to the proteostasis of disease-causing proteins. DnaJA1 and DnaJA2 are structurally similar JDPs that are related to prokaryotic *Escherichia coli* DnaJ and have been previously reported to be components of EVs. For example, DnaJA1 and DnaJA2 are components of mouse brain EVs (Brenna et al., 2020) while DnaJA1, DnaJA2 as well as DnaJC5 are components of EVs released from rat hippocampal neuronal cultures (Vilcaes et al., 2021). DnaJA1 associates with α-synuclein fibrils (Pemberton et al., 2011), modulates mutant huntingtin aggregation (Rodriguez-Gonzalez et al., 2020, Brehme et al., 2014), tau fibril clearance (Abisambra et al., 2012) and Abeta_42_ clearance (Brehme et al., 2014). DnaJA2 is a suppressor of tau aggregation *in vitro* and in cells (Mok et al., 2018, Irwin et al., 2021).

**Figure 3.**
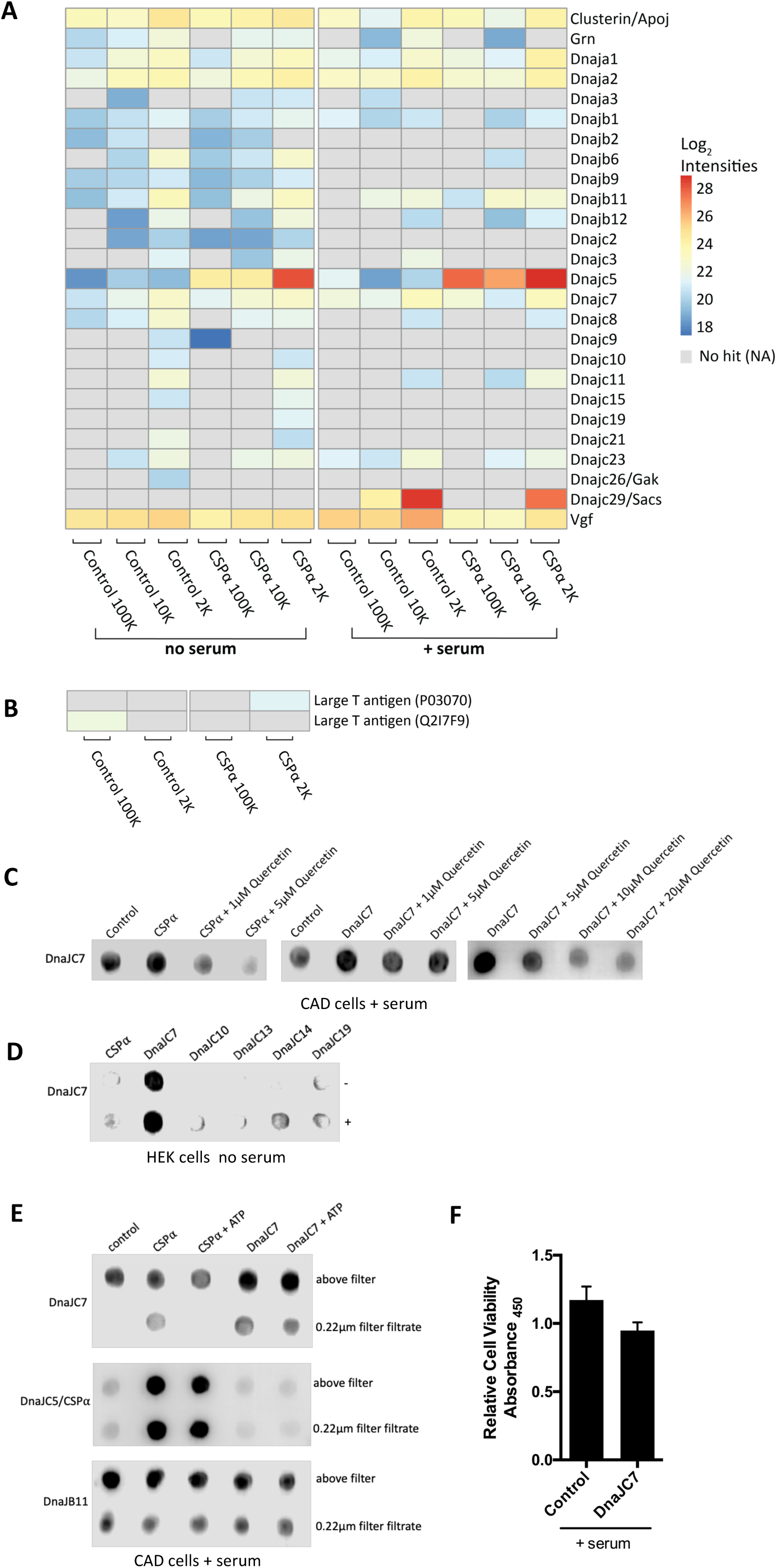
Identification of JDPs secreted from CAD cells. (A) Heat map of the exported JDPs identified +/- CSPα/DnaJC5 expression from cells grown in media without serum or 7.5% exosome-free FBS serum. Scale indicates intensity. (B) Heat map of exported SV40 large T antigen. (C) Dot blot of 10 µL of media collected from cells 48 hrs following CSPα/DnaJC5 or DnaJC7 expression treated with the indicated concentration of quercetin. (D) Media from HEK293 cells was collected 48 hrs following expression of the indicated JDPs in the absence (-) or presence (+) of GFP-72Q huntingtin^exon1^. (E) Media was further filtered through millex-GV 0.22µm PVDF filters in the presence or absence of 2mM ATP. Dot Blots are probed for DnaJC7, CSPα/DnaJC5 or DnaJB11 as indicated. (F) Cell viability.

DnaJC7 contains a C-terminal J domain and tetratricopeptide repeat (TPR) domains (Dilliott et al., 2022). The TRP domains are predicted to interact with Hsp90 and Hsp70 and may facilitate the flow of client proteins from Hsp70s to Hsp90s (Assimon et al., 2015). DnaJC7 associates with both tau and TDP-43 and is a suppressor of tau aggregation *in vitro* and in cells (Hou et al., 2021, Carrasco et al., 2023, Perez et al., 2023). Mutations in DnaJC7 cause dominantly inherited Amyotrophic Lateral Sclerosis (ALS) that typically does not feature tauopathy (Farhan et al., 2019). We verified the export of DnaJC7. Conditioned media was collected from CAD cells transiently expressing DnaJC7, filtered and 10 µL spotted on nitrocellulose and evaluated by dot blot. Endogenous DnaJC7 was found to be exported from control cells, from CSPα/DnaJC5-expressing cells and transient expression of DnaJC7 increased its secretion (Figure 3C, D, E). DnaJC7 expression reduced cell viability (Figure 3F). Quercetin, a natural compound that targets CSPα/DnaJC5 (Xu et al., 2010, Deruelle et al., 2021), was found to also reduce endogenous DnaJC7 secretion at low concentrations and transiently expressed DnaJC7 export at higher concentrations (Figure 3C). Conditioned media collected from HEK293 cells showed that DnaJC7 is exported in both the presence and absence of aggregation-prone proteins and confirmed that transient expression of DnaJC7 increased its secretion (Figure 3D). When media was filtered through 0.22 µm PVDF filters, DnaJC7 was detected in the filtrate as well as above the filter indicating it is associated with complexes of different sizes (Figure 3E). There was an inverse correlation between endogenous DnaJC7 in the filtrate and addition of 2mM ATP to the media following collection. ATP did not increase levels of DnaJC7, DnaJB11 and CSPα/DnaJC5 found in the filtrate.

Sacsin/DnaJC29 was notably abundant in the 2K pelleted fractions collected from cells grown in media containing exosome-depleted serum but not the 2K fractions collected from serum-starved cells (Figure 3A). Over 170 distinct mutations in sacsin/DnaJC19, a 520 kDa JDP, give rise to the childhood neurodegenerative disease, Autosomal Recessive Spastic Ataxia of Charlevoix-Saguenay (ARSACS) characterized by cerebellar atrophy (Bouchard et al., 1978). We therefore evaluated the media containing exosome-depleted serum (100 µg) for JDPs (data not shown). The Sacsin/DnaJC29 peptides, KVNALPEILKE and KFNGAQVNPKE, were identified. Sacsin/DnaJC29 was the only JDP detected in the media, indicating all other JDPs reported here originated from cells. Low levels of DnaJB1, known to associate with tau, TDP43, SOD1 and huntingtin, were found in most EVs (Irwin et al., 2021, Nachman et al., 2020, Carrasco et al., 2023, Ayala Mariscal et al., 2022, Serlidaki et al., 2020). In contrast, the classically secreted DnaJB11/ERdj3 was abundant in the 2K pellet as expected, but also detected in the high speed pelleted fractions (Figure 3A). Export of the nerve growth factor (VGF) and the chaperone, clusterin/apolipoprotein J, which are robustly secreted from CAD cells through the classical secretory pathway are shown for reference. Levels of the another cysteine rich lysosomal chaperone, progranulin (GRN), that is implicated in several neurodegenerative diseases, is also shown for reference (Rhinn et al., 2022, Braun, 2022, Kao et al., 2017). Clusterin and VGF were present in the 2K pellet, as anticipated, but also the 10K and 100K fractions. The presence of classically secreted chaperones and proteins in the 10K and 100K pellets suggest that they bind EVs as well as extracellular proteins.

In addition to eukaryotes, JDPs are found in bacteria and several viruses (Malinverni et al., 2023, Zhang et al., 2022). A notable feature of the pelleted EV fractions is the presence of large T antigen, a viral JDP essential for simian vacuolating virus 40 (SV40) DNA replication and transcriptional regulation (Mastrangelo et al., 1989, Sullivan et al., 2000). CAD cells are a mouse catecholaminergic cell line originally immortalized with polyomavirus SV40 (Qi et al., 1997). Low levels of large T antigen were detected in the 100K and 2K pellet fractions (Figure 3B).

In summary, CSPα/DnaJC5 increases the export of proteins, particularly in the 2K pellet and exported CSPα/DnaJC5 is found in a range of EVs separated by their pelleting properties. CSPα/DnaJC5 is not the only JDP that may influence the shuttling of proteins between cells. DnaJC7, DnaJA1 and DnaJA2 were major secreted JDPs in all conditions evaluated. Cells grown in the absence of serum released a greater number of distinct JDPs. We speculate that the distinct domains found in secreted JDPs may facilitate the secretion of diverse client proteins. Identifying the target proteins of exported JDPs requires further scrutiny and will need to be examined in a case-by-case manner.

To obtain a better understanding of the functional role of export, we evaluated the possible secretion of intracellular CSPα/DnaJC5-interacting proteins via the CSPα/DnaJC5-export pathway. CSPα/DnaJC5 is an established Hsc70/Hsp70 activator (Braun et al., 1996, Chamberlain and Burgoyne, 1997), that also associates with SGT (small glutamine-rich tetratricopeptide repeat-containing protein) (Tobaben et al., 2001) and Hsp90 (Sakisaka et al., 2002). In addition to assembly with other chaperones, several intracellular candidate substrates for CSPα/DnaJC5 have been proposed (Sharma et al., 2012, Sharma et al., 2011, Zhang et al., 2012, Leveque et al., 1998, Wu et al., 1999, Evans and Morgan, 2002, Magga et al., 2000, Sakisaka et al., 2002, Miller et al., 2003, Kyle et al., 2013, Lee et al., 2023). Seventeen out of 27 putative intracellular interacting proteins were found to be exported from both control and CSPα/DnaJC5-expressing cells. (Figure 4). As anticipated, Hsp90alpha, Hsp90beta and Hsc70/HSPA8 were abundant in EVs from control and CSPα/DnaJC5-expressing cells. Significant export of SLC3A2 and GTP binding proteins was found. SNAP23, a SNARE protein involved in EV secretion, is shown for reference (Liu et al., 2023). Association and disassociation with Hsp70s and client proteins are central to CSPα/DnaJC5’s intracellular activity, although extracellularly CSPα/DnaJC5 likely acts as a “holdase” meaning it may maintain association with client proteins (Braun, 2022).

**Figure 4.**
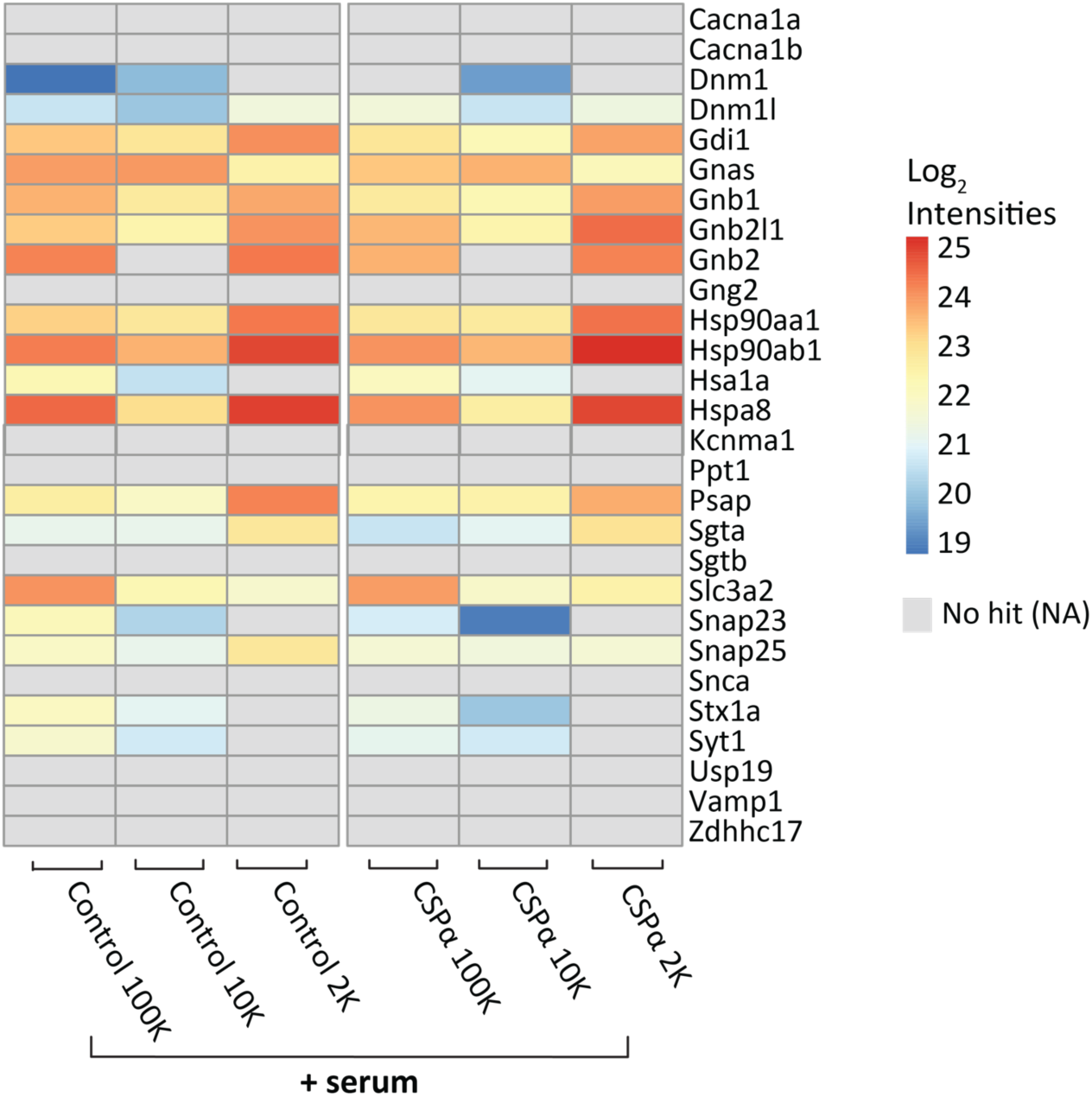
Export of predicted CSP⍺-interacting proteins. Heat map showing export of proteins reported to be CSPα/DnaJC5-interacting proteins. Scale indicates intensity.

We found 45 differentially exported as shown by volcano plots (Figure 5). Nineteen proteins showed at least a two fold upregulation in export and twenty six proteins were downregulated, indicating the impact of CSPα/DnaJC5 expression on protein export is broad (Figure 5). Among the most exported proteins were proteins known to function in DNA and RNA processing, as well as mitochondrial function, including the JDP DnaJC11, yet how they might contribute to synapse maintenance requires further study. A greater number of proteins were exported from serum-deprived cells expressing CSPα/DnaJC5. In CSPα/DnaJC5 null mice, multiple intracellular JDPs and other chaperones are downregulated suggesting a central role of CSPα/DnaJC5 in synaptic maintenance (Wang et al., 2023). We failed to see significant changes in the export of the reported CSPα/DnaJC5-interacting proteins with the exception of Hsp90 which had greater export in the 100K pellet of serum-deprived cells (Figure 4, 5). These results indicate that CSPα/DnaJC5 alters protein export in the absence of mutant-misfolded proteins.

**Figure 5.**
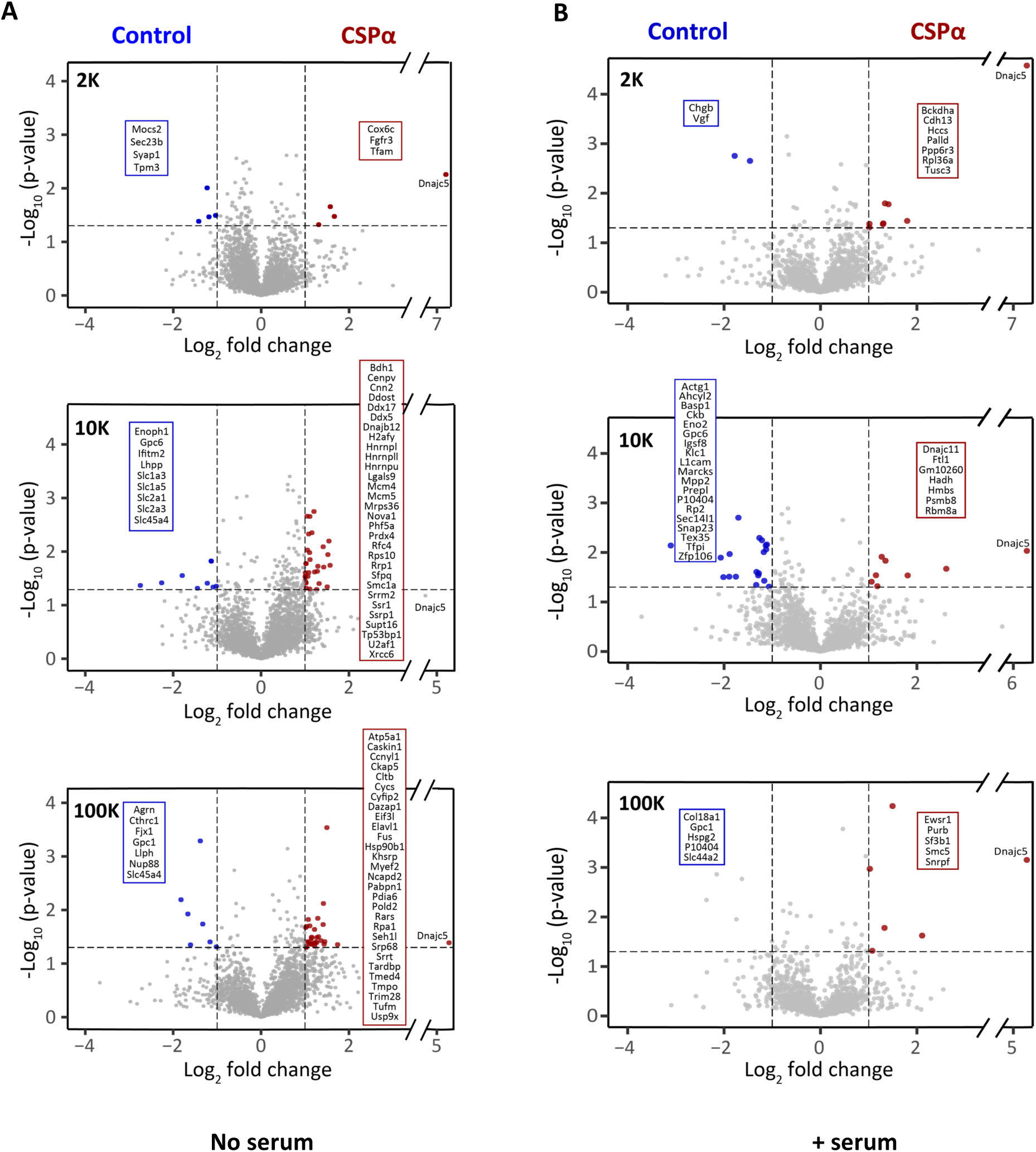
CSP⍺ drives changes in protein export. Volcano plot of proteins exported from control and CSPα/DnaJC5-expressing cells. (A) Serum starved cells. (B) Cells grown in 7.5% exosome-free FBS. Dotted vertical lines demarcate 2-fold changes and the dotted horizontal line represents. The difference between the CSPα/DnaJC5 and control secreted proteins was >1.3; *p* < 0.05), which represents a log2 fold change of 1 (for CSPα/DnaJC5) and -1 (for control). Each point represents a protein. Blue and red boxes represent significant changes and the UniProt names are indicated.

### Proteomic profiling of EVs secreted from cells expressing CSPα_L115R_ or CSPα_L116del_

Leu115Arg, Leu116del or C128Y mutations within the cysteine string region and duplication of a segment within the cysteine string region lead to adult-onset neuronal ceroid lipofusinosis (ANCL) a rapidly developing neurodegenerative lysosomal storage disease with no treatment (Benitez et al., 2011, Noskova et al., 2011, Velinov et al., 2012, Jedlickova et al., 2020, Huang et al., 2022). ANCL is one of 13 genetically distinct diseases called neuronal ceroid lipofuscinosis (CLNs) that are characterized by a progressive accumulation of highly autofluorescent lipofuscin deposits (Lee et al., 2022). Specifically, ANCL is characterized by ataxia, involuntary movements, seizures, cognitive deterioration and patients usually die within 10 years of diagnosis (Lee et al., 2022). The most extensively studied Leu115Arg and Leu116del mutants have altered palmitoylation and localization and show aggregation but the the molecular mechanisms by which the mutant versions of CSPα/DnaJC5 cause ANCL are not understood in detail (Zhang and Chandra, 2014, Greaves et al., 2012, Henderson et al., 2016, Imler et al., 2019, Diez-Ardanuy et al., 2017, Naseri et al., 2020, Lee et al., 2023, Lopez-Begines et al., 2023). Mice expressing CSPα_L115R_ or CSPα_L116del_ do not have a reduced life span but show pathological lipofuscin deposits, with greater lipofuscinousis observed in mice with the L115R mutation (Lopez-Begines et al., 2023). These reports suggest that Leu115Arg or Leu116del mutants may have have pathological interactions with yet unknown proteins thereby causing a toxic gain-of-function leading to lipofuscinosis and neurodegeneration.

To gain further insight into the molecular deficiencies in ANCL, we inspected the exported proteome from cells expressing CSPα_L115R_ or CSPα_L116del_. While Leu115Arg and Leu116del mutations within the cysteine string region of CSPα/DnaJC5 cause a pathogenic progression of events leading to lipofusciosis, they do not alter association with Hsp70s (Zhang and Chandra, 2014). Autofluorescence analysis was conducted on the pelleted fractions (n = 19) prepared from cells grown in medium supplemented with 7.5% exosome-depleted serum (Supplementary Figure 1). Overall autofluorescent intensity was higher in the 2K pellets from CSPα, CSPα_L115R_ and CSPα_L116del_ expressing cells_._ The emission spectrum of the autofluorescence showed a similar profile in all 2K pellets, that is, low blue, high green, medium red and low far red. We failed to see higher autofluorescence in EVs from cells expressing mutants compared to wild type CSPα/DnaJC5-expressing cells, although the 10K pellet from CSPα_L115R_-expressing cells and the 100K pellet from CSPα_L116del-_expressing cells showed a narrowing of the spectrum, (i.e. more green and red). How EV autofluorescence connects to CLNs is not known. In mice, the absence or overexpression of wild-type CSPα/DnaJC5 does not cause lipofuscin deposits and determining why mutations in CSPα/DnaJC5 present as lipofuscin deposits or if EVs are involved requires further investigation.

There are constrasting observations resulting from different techniques and assays used to study export of misfolded proteins by CSPα_L115R_, CSPα_L116del_. We have reported that CSPα_L115R_, CSPα_L116del_ export GFP-72Q huntingtin^exon1^ and SOD-1^G93A^ from CAD cells (Deng et al., 2017). Lee and colleagues have found that CSPα_L115R_, CSPα_L116del_ do not export GFP1-10 from U2OS cells (Lee et al., 2023) and Wu and colleagues report that CSPα_L115R_, CSPα_L116del_ do not export α-synuclein from HEK293 cells (Wu et al., 2023). Although, unfractionated samples from CAD cells contained less CSPα_L115R_ and CSPα_L116del_ export relative to wild type (Figure 1), the low speed (2K) pellets show more CSPα_L115R_ or CSPα_L116del_ (Figure 6). Twenty two proteins released from CSPα_L115R_ - and 11 proteins released from CSPα_L116del_ -expressing cells were significantly altered compared to CSPα/DnaJC5-expressing cells (Figure 6A, B). Export of the ribosomal proteins, Pelo and Eiflax, from both CSPα_L115R_ and CSPα_L116del_ expressing cells was reduced, but other changes in the protein export were unique to specific mutations. Proteins that demonstrated increased export in the presence of mutants included proteins involved in cholesterol transport, transcription and cytoskeleton remodeling (Figure 6). These results show that CSPα_L115R_ and CSPα_L116del_ alter the export of proteins and suggest that changes in the export of proteins may underlie the pathogenesis of these mutants.

**Figure 6.**
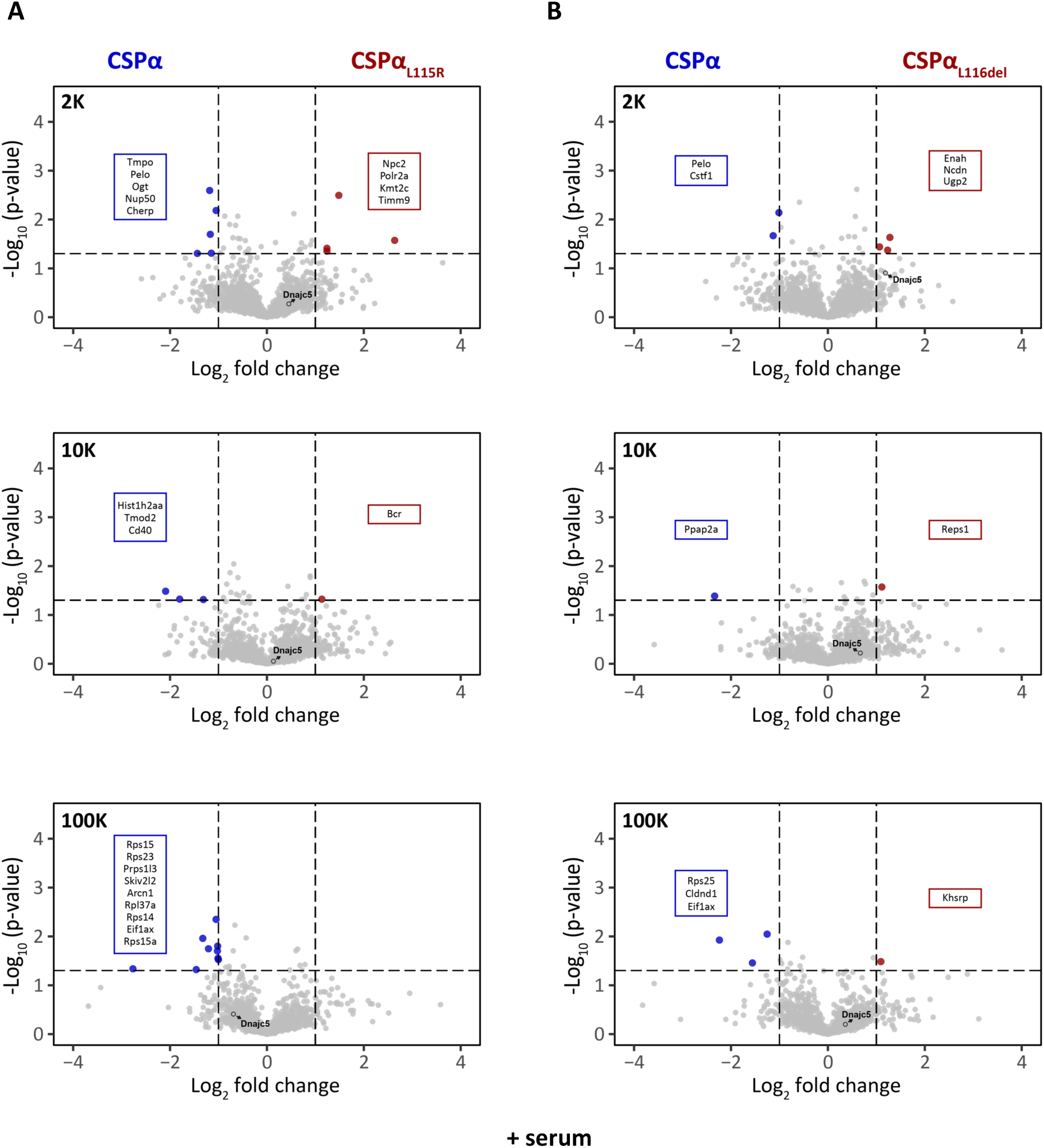
CSPα_L115R_ and CSPα_L116del_ mediate changes in protein export. Volcano plot of proteins exported from (A) CSPα/DnaJC5 and CSPα_L115R_, (B) CSPα/DnaJC5 and CSPα_L116del_ -expressing cells. (C) Metascape pathway analysis for CSPα/DnaJC5 vs control-no serum, CSPα/DnaJC5 vs control-serum, CSPα/DnaJC5 vs CSPα_L115R_ and CSPα/DnaJC5 vs CSPα_L116del._ Blue and red boxes represent significant changes and the UniProt names are indicated.

To gain further insight into the role of CSPα/DnaJC5 export we examined the export of endogenous α-synuclein, amyloid beta, APOE, PARK7, prion protein, huntingtin, SOD-1, and TDP-43. Misfolded versions of these proteins, are associated with various neurodegenerative diseases, as described earlier, and misfolded α-synuclein, huntingtin, SOD-1, and TDP-43 are associated with CSPα/DnaJC5 export (Wu et al., 2023, Deng et al., 2017, Fontaine et al., 2016, Lee et al., 2018, Pink et al., 2021). Intriguingly, α-synuclein overexpression can rescue the neurodegeneration associated with CSPα/DnaJC5 knockout (Chandra et al., 2005). Wild-type α- synuclein, tau/MAPT, huntingtin and APOE were not found in the pelleted fractions obtained from media collected from control cells or cells expressing CSPα/DnaJC5 or CSP mutants, when grown in the serum (Figure 7). Serum starved cells exported low levels of tau/MAPT and huntingtin. We did not detect evidence of increased export of these proteins between control and CSPα/DnaJC5-expressing cells or major differences in the overall pattern of export from CSPα/DnaJC5 and CSPα/DnaJC5 mutant-expressing cells. Nevertheless, post translational modifications of exported proteins is implicated in multiple neurodegenerative contexts and remains to be explored despite the similar protein export signatures.

**Figure 7.**
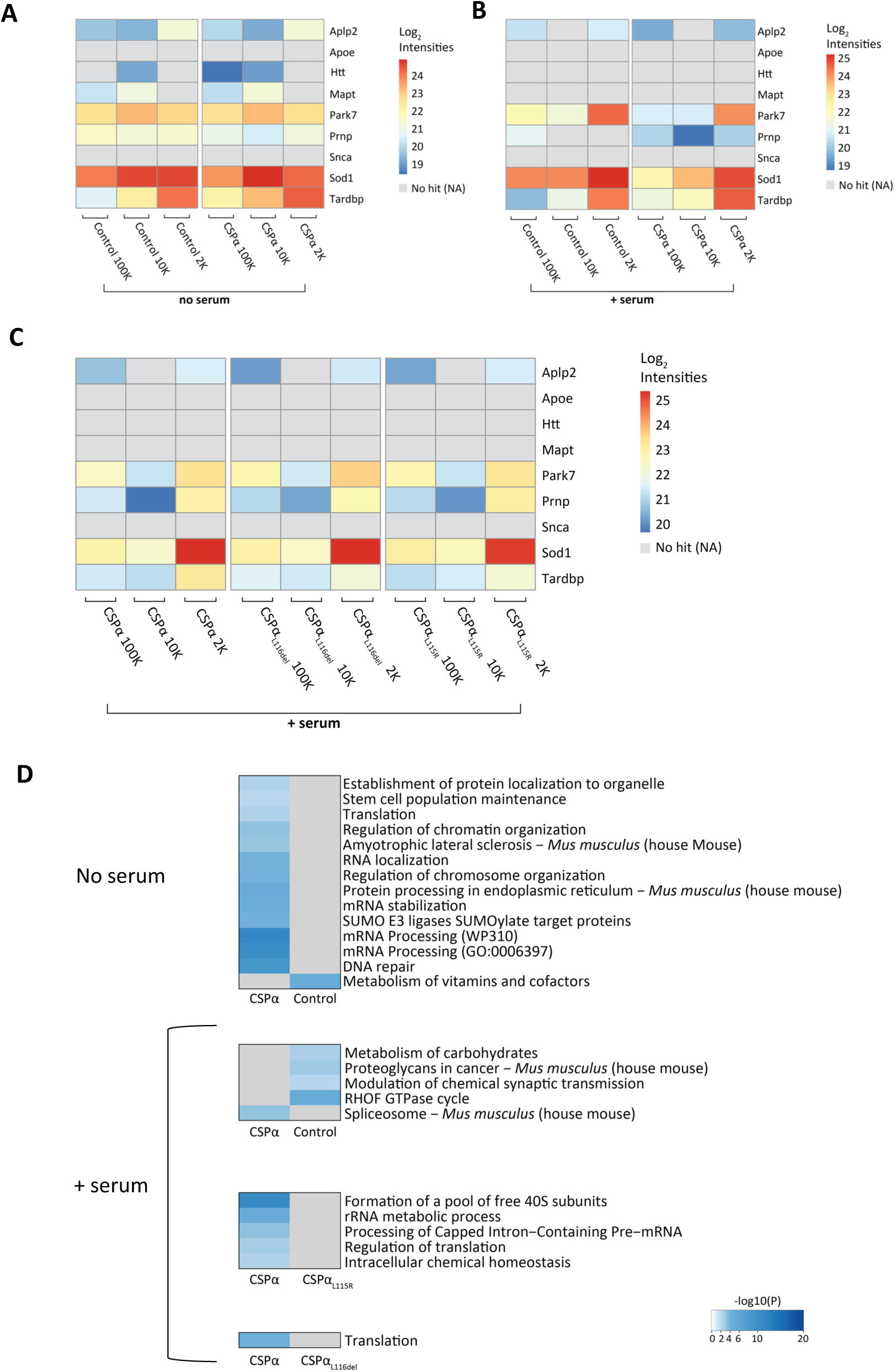
Export of SOD1 and TDP43 from cells expressing CSPα/DnaJC5 or CSPα/DnaJC5 mutants. Heat map of exported wild-type proteins. The proteins shown are implicated in neurodegenerative diseases when misfolded. (A) Serum starved cells. (B, C) Cells grown in 7.5% exosome-free FBS. A, B, C show three separate comparisons as the data was analyzed independently. Scale indicates intensity. The heat map shows proteins that were independently identified in 2 (out of 3) biological samples. (D) The top pathways identified based on significantly changed proteins.

Finally, we combined the significant proteomic hits from the three pellets for pathway analysis using metascape (Zhou et al., 2019). Identified pathways based on curated protein-protein association are shown in Figure 7D. Pathway analysis revealed unanticipated differences in CSPα/DnaJC5, CSPα_L115R_ and CSPα_L116del_ protein exprot, despite the absence of changes in export of endogenous α-synuclein, amyloid beta, APOE, PARK7, huntingtin, SOD-1, and TDP-43 suggesting additional pathways need be considered in ANCL pathogenesis. More work is needed to fully understand the effects of CSPα/DnaJC5 mutations on different export targets and the downstream effects of these changes.

### Conclusions

Identification of proteins exported from CSPα/DnaJC5 expressing cells provides a rational basis for investigating the role that CSPα/DnaJC5 plays in protein transit through the central nervous system and the mechanisms by which CSPα/DnaJC5 maintains CNS proteostasis. The work presented herein systematically and comprehensively validates proteins exported in EVs from CSPα/DnaJC5, CSPα_L115R_ and CSPα_L116del_ -expressing cells. This study supports a role for CSPα/DnaJC5 in intercellular communication. CSPα/DnaJC5 increases the export of proteins, particularly in the 2K pellet and CSPα/DnaJC5 is found in a range of EVs separated by their pelleting properities. Unbiased proteomic analysis revealed both upregulation and downregulation of protein export in CSPα/DnaJC5-expressing cells and that export signatures of CSPα_L115R_ and CSPα_L116del_ expressing cells are distinct. Our screen greatly expands the repertoire of JDPs known to be exported. A role for CSPα/DnaJC5 in cell-to-cell communication aligns with work from others showing the contribution of CSPα/DnaJC5 in the modulation of neuron-glia interactions (Wang et al., 2023) and cell-cycle progression of neural progenitor cells (Nieto-Gonzalez et al., 2019).

CSPα/DnaJC5 is central to synaptic integrity and CSPα/DnaJC5 dysfunction a common molecular thread across several neurodegenerative disorders including; the lysosomal storage disorder adult-onset neuronal ceroid lipofusinosis (ANCL) (Benitez et al., 2011, Noskova et al., 2011, Velinov et al., 2012, Jedlickova et al., 2020, Huang et al., 2022, Donnelier et al., 2015) (ANCL), Alzheimer’s disease (Zhang et al., 2012, Tiwari et al., 2015, Rupawala et al., 2022), Parkinson’s disease (Chandra et al., 2005, Wu et al., 2023, Calo et al., 2021, Guo et al., 2023, Lee et al., 2018) and Huntington’s disease (Deng et al., 2017, Pink et al., 2021). The *in vivo* propagation of toxic-mutant proteins between contiguous neuroanatomical regions in the CNS, aligns with CSPα/DnaJC5 ejection of mutant α-synuclein, TDP-43, tau, SOD-1 and huntingtin from cells and (Wu et al., 2023, Deng et al., 2017, Fontaine et al., 2016, Lee et al., 2018, Pink et al., 2021, Graykowski et al., 2020). Our results add to this series of observations by identifying a set of native proteins exported from CSPα/DnaJC5-expressing cells in the absence of mutant, oligomeric disease-causing proteins.

### Limitations of the Study

The CSPα/DnaJC5 exported proteome reported in the work was detected in CAD cells under no serum (stressed) or 7.5% exosome-depleted serum (non-stressful) conditions. The CSPα/DnaJC5 exported proteome will be different under other proteomic stress conditions and most likely vary amoung cell types. Degeneration in the absence of CSPα/DnaJC5 is activity-dependent (Garcia-Junco-Clemente et al., 2010, Schmitz et al., 2006) suggesting that neuronal activity may be coupled to CSPα/DnaJC5 export, something that is not tested in this report. In addition, during aging proteostasis declines (Molzahn et al., 2023), age-dependent changes in CSPα/DnaJC5 export is not tested in this report. As with any proteomic study, a fraction of proteins were likely not identified and future studies may identify other exported proteins. In this light, our work provides a starting point to begin to understand how the CSPα/DnaJC5 and other JDP export pathways work and how they may be targeted for future pharmacological interventions. Finally, it will be necessary to assess the physiological uptake pathways for CSPα/DnaJC5 exported cargo to understand the mechanisms underlying synapse maintenance.

## Methods

### Key resource table

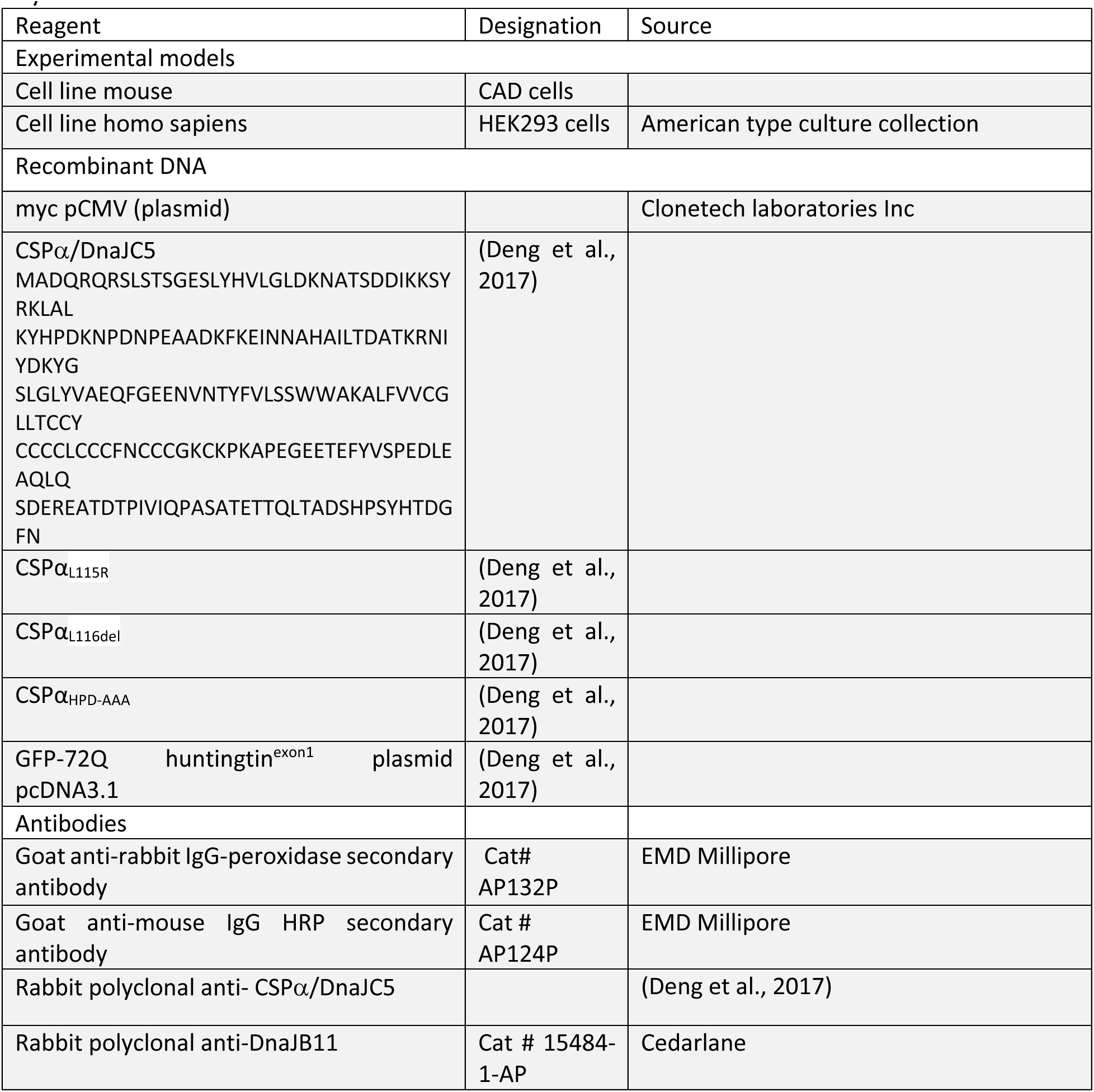
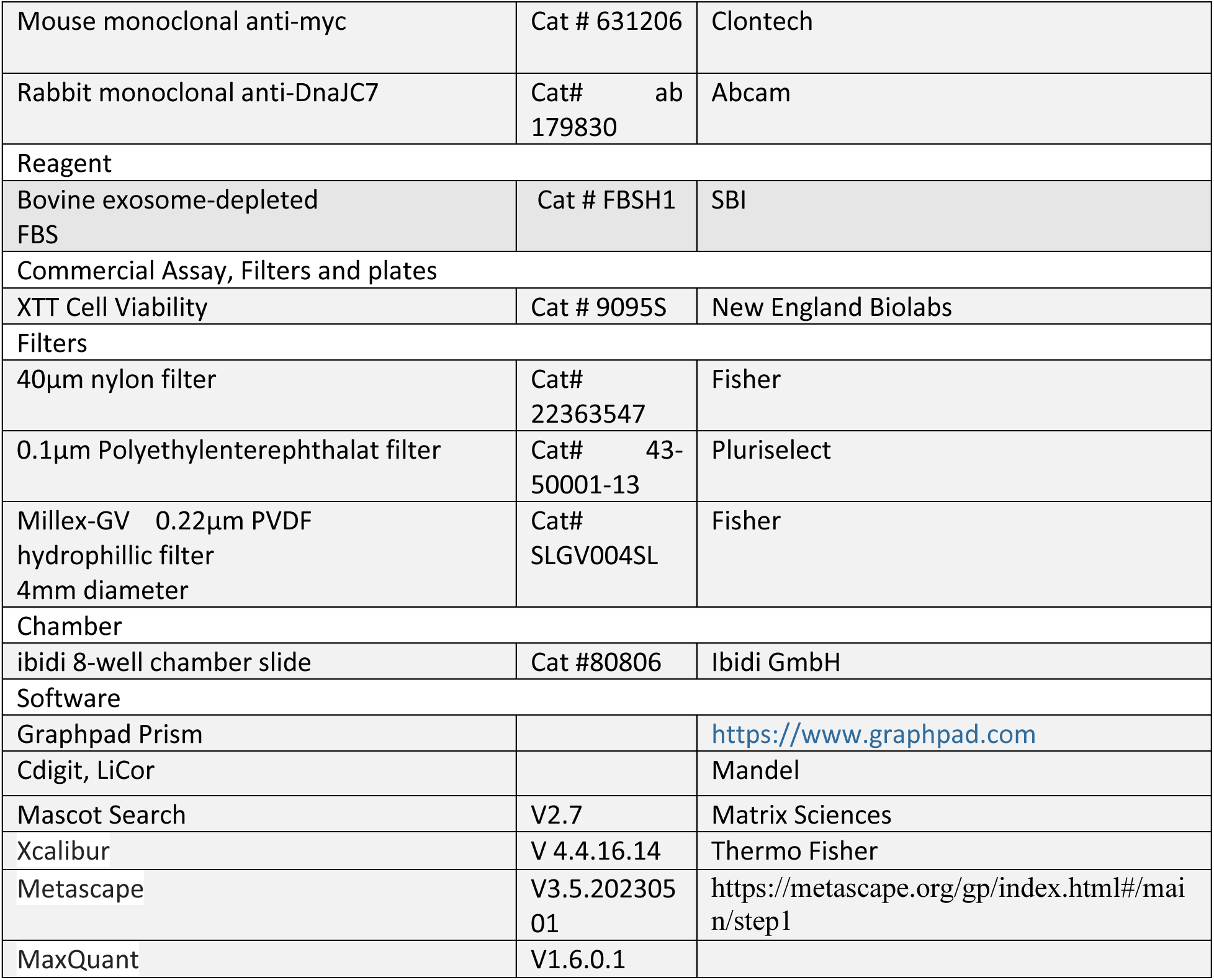

### Cell Culture and Extracellular Vesicles

CAD cells (catecholaminergic neuronal cells) or HEK293T cells were seeded in 10 cm culture dishes in Dulbecco’s Modified Eagles’s Medium (DMEM-F12), supplemented with 10% Fetal Bovine Serum, 1% penicillin and 100µg/ml streptomycin and maintained at 37 °C, 5% CO_2_ atmosphere. Cells were transfected with plasmids encoding different constructs of CSPα and GFP-72Q huntingtin^exon1^ using Lipofectamine 3000 reagent in OptiMEM medium. At 6 h after transfection, cell culture medium was replaced with either serum free or 7.5% exosome depleted serum (validation) supplemented Dulbecco’s Modified Eagle’s Medium (+Lglutamine, +15mm HEPES, - phenol red). Conditioned media was collected at 24 h or 48 h after transfection and cellular debris were removed by filtration through 40 µm (nylon) and 1 µm (PET; polyethylentererephthalat) filters (pluriSelect). Negative pressure via a syringe was applied to the 1µm filter. Media was then centrifuged at 2,000 x *g* for 20 min at 4 °C (2K pellet). The supernatant was transferred to new tubes and centrifuged for 40 min at 10,000 x *g* (10K pellet) and finally for 90 min at 100,000 x *g* (100K pellet). The 2K pellet was resuspended in PBS. The 10K and 100K pellets were washed in sterile PBS, recentrifuged and resuspended in PBS. Protein concentration was measured using Nanodrop (Fisher Scientific). Following media collection, cell viability was determined utilizing an XTT assay (New England Biolabs). 10 µL of unfractionated conditioned media or 5 µL of the 2K, 10K and 100K pellets were spotted on nitrocellulose membrane (0.2 µm pore size). Membranes were blocked in tris-buffered saline containing 0.1% Tween 20, 1% BSA and then incubated with primary antibody overnight at 4 °C. The membranes were washed and incubated with horseradish peroxidase-coupled secondary antibody for 2 h at room temperature. Bound antibodies on the membrane were detected by incubation with Pierce chemiluminescent reagent and exposure to Cdigit, LiCor (Mandel). Where indicated media was applied to Millex 0.22μm PVDF hydrophillic filter for further fractionation (overnight, 4 °C, 1 x *g*).

### Electron Microscopy

Electron microscopy was performed on pellets stored at −80 °C and never unfrozen. EV suspension in PBS was deposited carbon-coated 200 mesh copper grids that were glow charged for 1 min. Samples were washed with ammonium acetate (0.1 M and 0.01 M), pH 7.4 and stained with 2% (w/v) uranyl acetate and the grids were air-dried. The samples were viewed with a FEI Tecnai F20 transmission electron microscope under an acceleration voltage of 200 kV and electron micrographs were recorded on an Eagle 4 k × 4 k CCD camera.

### Autofluorescence Microscopy

Autofluorescence imaging was performed on a Nikon Inverted Ti2 A1R confocal microscope (Nikon Instruments Inc., Melville, USA) using a 60x 1.4NA oil lens. Samples were pipetted directly into an ibidi 8-well chamber slide (Cat.#80806, Ibidi GmbH, Gräfelfing, Germany) and allowed to settle for 30 minutes. Signals were collected in four channels with the following bandpass filters: 450/50 (Blue), 525/50 (Green), 595/50 (Red), and 700/75 (Far Red). The sample was excited with four laser lines, 405, 488, 561, and 638nm corresponding to the four collection channels, with all four laser lines set to 1% power for comparable excitation levels.

The images were analyzed in ImageJ by applying an automated background removal threshold (Triangle threshold) to avoid sample bias. After thresholding, the average image intensity was calculated from the foreground pixels. Thresholding and image intensity calculations were applied separately to each color channel in each image to account for the differences in signal level between channels. Standard error was calculated from the standard deviation of the pixel intensities for each group for graphing purposes.

### Proteomics and Mass Spectrometry Analysis

Proteomics analyses were performed and analyzed at the Southern Alberta Mass Spectrometry Facility (SAMS), University of Calgary. Three independent sets of 2K, 10K and 100K pellets from serum free media and three independent sets of 2K, 10K and 100K pellets from 7.5% exosome depleted serum (validation) were analyzed. 100 µg of each protein sample was subjected to TCA precipitation (final concentration 20%) on ice for 30 min, centrifuged at 17,000 x *g* for 15 min and washed 3 times with cold acetone prior to air drying. Dried proteins were resolubilized with 8 M urea in 100 mM Tris-HCl pH 8.0. The cysteines were reduced with 10 mM Dithiothreitol (DTT) at 37°C for 30 min and alkylated with iodoacetamide (IAA) at room temperature for 20 min. Reduced and alkylated proteins were then digested with trypsin overnight at 37 °C. Samples were freeze dried and desalted on UPLE reverse phase columns.

Tryptic peptides were analyzed on an Orbitrap Fusion Lumos Tribrid mass spectrometer (Thermo Scientific) operated with Xcalibur (version 4.4.16.14) and coupled to a Thermo Scientific Easy-nLC (nanoflow Liquid Chromatography) 1200 system. Tryptic peptides were loaded onto a C18 trap (75 um x 2 cm; Acclaim PepMap 100, P/N 164946; ThermoScientific) at a flow rate of 2 µL/min of solvent A (0.1% formic acid in LC-MS grade water). Peptides were eluted using a 120 min gradient from 5 to 40% (5% to 28% in 105 min followed by an increase to 40% B in 15 min) of solvent B (0.1% formic acid in 80% LC-MS grade acetonitrile) at a flow rate of 0.3 μL/min and separated on a C18 analytical column (75 um x 50 cm; PepMap RSLC C18; P/N ES803; ThermoScientific). Peptides were then electrosprayed using 2.1 kV voltage into the ion transfer tube (300 °C) of the Orbitrap Lumos operating in positive mode. The Orbitrap first performed a full MS scan at a resolution of 120,000 FWHM to detect the precursor ion having a *m*/*z* between 375 and 1,575 and a +2 to +7 charge. The Orbitrap AGC (Auto Gain Control) and the maximum injection time were set at 4 x 10^5^ and 50 ms, respectively. The Orbitrap was operated using the top speed mode with a 3 sec cycle time for precursor selection. The most intense precursor ions presenting a peptidic isotopic profile and having an intensity threshold of at least 5000 were isolated using the quadrupole and fragmented with HCD (30% collision energy) in the ion routing multipole. The fragment ions (MS^2^) were analyzed in the ion trap at a rapid scan rate. The AGC and the maximum injection time were set at 1 x 10^4^ and 35 ms, respectively, for the ion trap. Dynamic exclusion was enabled for 45 sec to avoid of the acquisition of same precursor ion having a similar m/z (plus or minus 10 ppm).

For identification, the Lumos raw data files (*.raw) were converted into MaxQuant and the data searched against the UniProt mouse data base and, where indicated, viral database. Mascot Generic Format (MGF) using RawConverter (v1.1.0.18; The Scripps Research Institute) operating in a data dependent mode. Monoisotopic precursors having a charge state of +2 to +7 were selected for conversion. This mgf file was used to search a Mus musculus database using Mascot algorithm (Matrix Sciences; version 2.7). Search parameters for MS data included trypsin as enzyme, a maximum number of missed cleavage of 1, a peptide charge equal to 2 or higher, cysteine carbamidomethylation as fixed modification, methionine oxidation as variable modification and a mass error tolerance of 10 ppm. A mass error tolerance of 0.6 Da was selected for the fragment ions. Only peptides identified with a score having a confidence higher than 95% were kept for further analysis. The Mascot data files were imported into Scaffold (v4.3.4, Proteome Software Inc) for comparison of different samples based on their mass spectral counting.

Spectral data were matched to peptide sequences in the human UniProt protein database using the Andromeda algorithm (Cox et al., 2011) as implemented in the MaxQuant (Cox and Mann, 2008) software package v.1.6.10.23, at a peptide-spectrum match FDR<0.01. Search parameters included a mass tolerance of 20 p.p.m. for the parent ion, 0.5 Da for the fragment ion, carbamidomethylation of cysteine residues (+57.021464 Da), variable N-terminal modification by acetylation (+42.010565 Da), and variable methionine oxidation (+15.994915 Da). N-terminal and lysine heavy (+34.063116 Da) and light (+28.031300 Da) dimethylation were defined as labels for relative quantification. The cleavage site specificity was set to Trypsin/p for the proteomics data, with up to two missed cleavages allowed. Significant outlier cut-off values were determined after log (2) transformation by boxplot-and-whiskers analysis using the BoxPlotR tool (Spitzer et al., 2014). The data were deposited into ProteomeXchange via the PRIDE database and are freely available: PXD046707.

To identify protein–protein interactions, the STRING (Search Tool for the Retrieval of Interacting Genes) database was used to illustrate interconnectivity among proteins. Protein-protein interactions relationship were encoded into networks in the STRING v11 database (https://string-db. org). Data were analyzed using the homo sapiens as the organism (FDR=0.05). Metascape analysis was also used to identify enriched pathways. Protein-protein interactions relationship were encoded into networks using the Metascape website (https://metascape.org/), and the enriched pathways were plotted as heatmaps.

## Funder

Natural Science and Research Council of Canada RGPIN/04826-2017

The funders had no role in study design, data collection and interpretation.

---

**Supplementary Figure 1.**
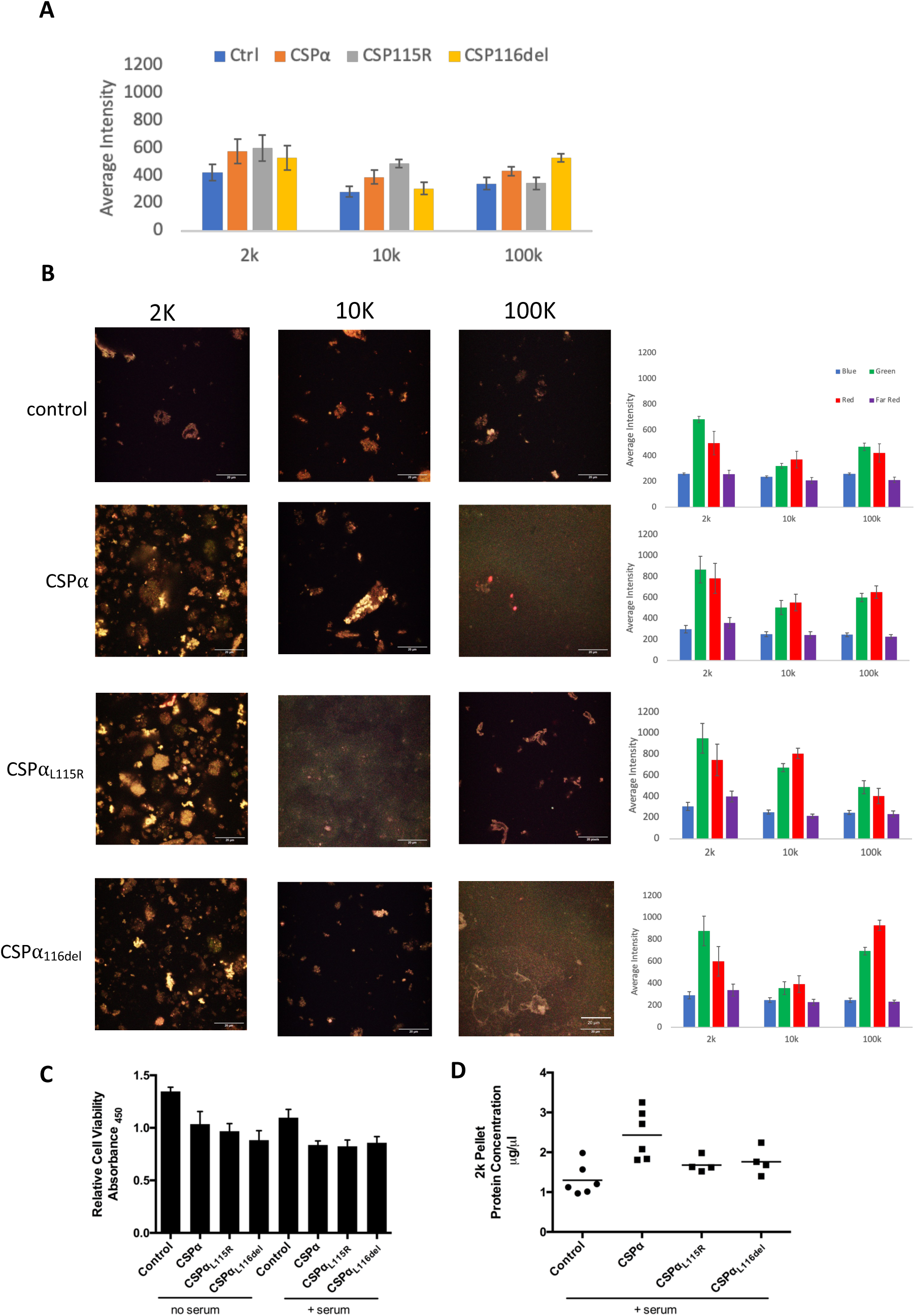
Autofluoresence of EVs released from control, CSPα, CSPα_L115R_ and CSPα_L116del_-expressing cells. (A) Overall autofluorescent intensity in the 2K, 10K and 100K pellets prepared from media that was collected from control cells or cells expressing CSPα, CSPα_L115R_, or CSPα_L116del_ 48 h after transfection. Each pellet was imaged 19 times. (B) Representative autofluorescent images. (C) Cell viability (D) protein content of the 2K pellets.

## Notes

### Competing Interest Statement

The authors have declared no competing interest.

